# Behavioral and Functional Profiling of *Acomys cahirinus* Fibroblasts Reveals Enhanced Matrix Remodeling Capacity

**DOI:** 10.64898/2026.07.07.737114

**Authors:** Nico Macaluso, Meera Bhat, Angela Lu, Yuanhui Cheng, Long T. Nguyen, Piyush K. Jain, Jude M. Phillip

## Abstract

The African spiny mouse (*Acomys cahirinus*) exhibits a unique capacity among mammals for scarless tissue regeneration, making it a compelling model for investigating the cellular mechanisms underlying regenerative healing. To determine how cellular heterogeneity and specific phenotypes influence fibroblast behavior, we established an immortalized *Acomys* fibroblast line along with a CRISPR/Cas9-mediated Col3A1 knockout variant and a DNA damage-induced senescent population. Compared with *Mus musculus,* NIH 3T3 fibroblasts, *Acomys* cells displayed distinct morphology, similar migration speeds, reduced directional persistence, and greater biophysical heterogeneity. While previous studies have linked regenerative wound healing to the elevated expression of collagen type III (Col3A1), CRISPR-mediated knockout of Col3A1 in *Acomys* fibroblasts yielded comparable biophysical profiles to wild-type cells in 2D culture. To examine additional contributors to the enhanced wound-like matrix environment, we established a senescence model in which *Acomys* fibroblasts exhibited elevated resistance to DNA-damaging agents, complete loss of proliferation, and altered single-cell morphology. In 3D collagen gel contraction assays, Col3A1 knockout attenuated matrix remodeling capacity, whereas the introduction of a small fraction of senescent cells enhanced gel contraction and remodeling dynamics, suggesting that senescent fibroblasts can modulate collective matrix behaviors. Together, these findings demonstrate that both Col3A1 expression and senescence-associated cell states contribute to fibroblast-driven matrix remodeling, highlighting *Acomys* fibroblasts as a valuable model for investigating how cellular heterogeneity and senescence-associated cell phenotypes could influence regenerative wound healing.

## INTRODUCTION

Tissue regeneration in adult mammals is typically limited, often resulting in fibrotic scarring rather than complete restoration of original tissue architecture (Gaire et al., 2021). In contrast, the African spiny mouse *Acomys cahirinus* (*Acomys*) exhibits a remarkable ability to regenerate complex tissues, including skin, hair follicles, and cartilage, without scarring (Gaire et al., 2021; Seifert et al., 2012). This distinctive regenerative capacity positions *Acomys* as a compelling model to study the mechanisms of scarless wound healing. However, the cellular phenotypes underlying this process, particularly compared with canonical mammalian systems such as *Mus musculus* (*Mus*), remain poorly understood. Deeper characterization of these phenotypes, especially in controlled *ex vivo* systems, could yield functional insights that inform next-generation therapies.

Fibroblasts are central mediators of wound healing, orchestrating extracellular matrix (ECM) deposition, remodeling, and resolution of inflammation (Cialdai et al., 2022; Tracy et al., 2016). In typical mammalian wound repair, fibroblast activation leads to myofibroblast differentiation and subsequent scar formation (Li & Wang, 2011). By contrast, *Acomys* fibroblasts contribute to regeneration without progressing to fibrosis, suggesting intrinsic differences in their functional properties (Gaire et al., 2021; Seifert et al., 2012). However, the cellular behaviors and molecular mechanisms enabling this regenerative phenotype are incompletely understood. A distinguishing feature of *Acomys* wound repair is its ECM composition. While typical mammalian wound repair results in dense, highly aligned type-1 collagen-rich scar tissue, regenerative healing in *Acomys* is characterized by a more fetal-like ECM with elevated type-III collagens and reduced fibrosis (Brant et al., 2015; Stewart et al., 2025).

Type-III collagen, encoded by the Col3A1 gene, plays a critical role in maintaining tissue integrity, regulating matrix organization, and modulating the cellular response to injury (Kuivaniemi & Tromp, 2019; Omar et al., 2021). This collagen subtype is abundant during early wound repair where it contributes to a compliant matrix that supports tissue remodeling before being replaced by type-I collagens during scar maturation. Notably, *Acomys* wounds exhibit elevated and sustained Col3A1 expression relative to scar-forming *Mus* wounds, with recent work demonstrating that type-III collagens directly influence matrix architecture, fibroblast activation, and mechano-sensing during tissue repair (Brant et al., 2015; Stewart et al., 2025). Conversely, reduced type-III collagens are associated with increased fibrosis, aberrant collagen alignment, impaired wound healing, and connective tissue disorders such as vascular Ehlers-Danlos syndrome (Kuivaniemi & Tromp, 2019; Omar et al., 2021). These findings suggest that type-III collagens may function not only as a marker of regenerative healing but as an active regulator of fibroblast behavior, prompting the investigation of *Acomys* fibroblasts with genetic deletion of Col3A1.

Cellular senescence is a state of irreversible growth arrest that plays a complex role in tissue repair. Transient senescence can facilitate wound healing through the senescence-associated secretory phenotype (SASP), which is the coordinated release of signaling molecules that promote tissue remodeling. However, chronic senescence, for instance during aging (Childs et al., 2015), contributes to tissue dysfunction and fibrosis (Saito et al., 2024; Wilkinson & Hardman, 2020, 2022). Understanding how *Acomys* fibroblasts navigate senescence pathways may reveal strategies to promote regenerative healing while mitigating fibrotic outcomes. Although senescence likely plays a central role in wound repair in *Acomys* tissues, induction of senescence in *Acomys* cells within *ex vivo* cultures yields mixed outcomes. One study inducing senescence in *Acomys* cells using hydrogen peroxide (via reactive oxidative stress, ROS) showed that *Acomys* cells were resistant to senescent induction. However, it is unclear whether this resistance to senescence induction was related to the method of induction or whether it was an artifact of the *in vitro* culture (Saxena et al., 2019).

Here, we addressed these gaps by establishing an immortalized *Acomys* fibroblast model, and using *ex vivo* approaches, we systematically characterized fibroblast phenotypes. We also generated a CRISPR/Cas9-mediated Col3A1 knockout line. By comparing the single-cell behaviors of these cells with *Mus* fibroblasts, we investigated morphological, proliferative, and functional characteristics under basal conditions and following senescence induction. These findings provide insights into the cellular determinants of scarless wound healing and the putative roles of Col3A1 and senescence in modulating fibroblast function.

## RESULTS

### *Acomys* fibroblasts exhibit distinct biophysical behaviors relative to *Mus* fibroblasts

The diverse cell populations within the dermal environment likely contribute to the unique regenerative ability of *Acomys*. Skin fibroblasts play a central role in this process (Talbott et al., 2022) and their biophysical properties not only govern function but also classify and predict phenotypic outcomes (Kamat et al., 2024). To better understand the factors contributing scarless healing, we obtained an immortalized *Acomys* fibroblast line and compared its two-dimensional (2D) biophysical behaviors to those of *Mus* NIH 3T3 fibroblasts. Both cellular and nuclear parameters were analyzed to quantify fibroblast phenotypes (**Figure 1A-B**)

**Figure 1.**
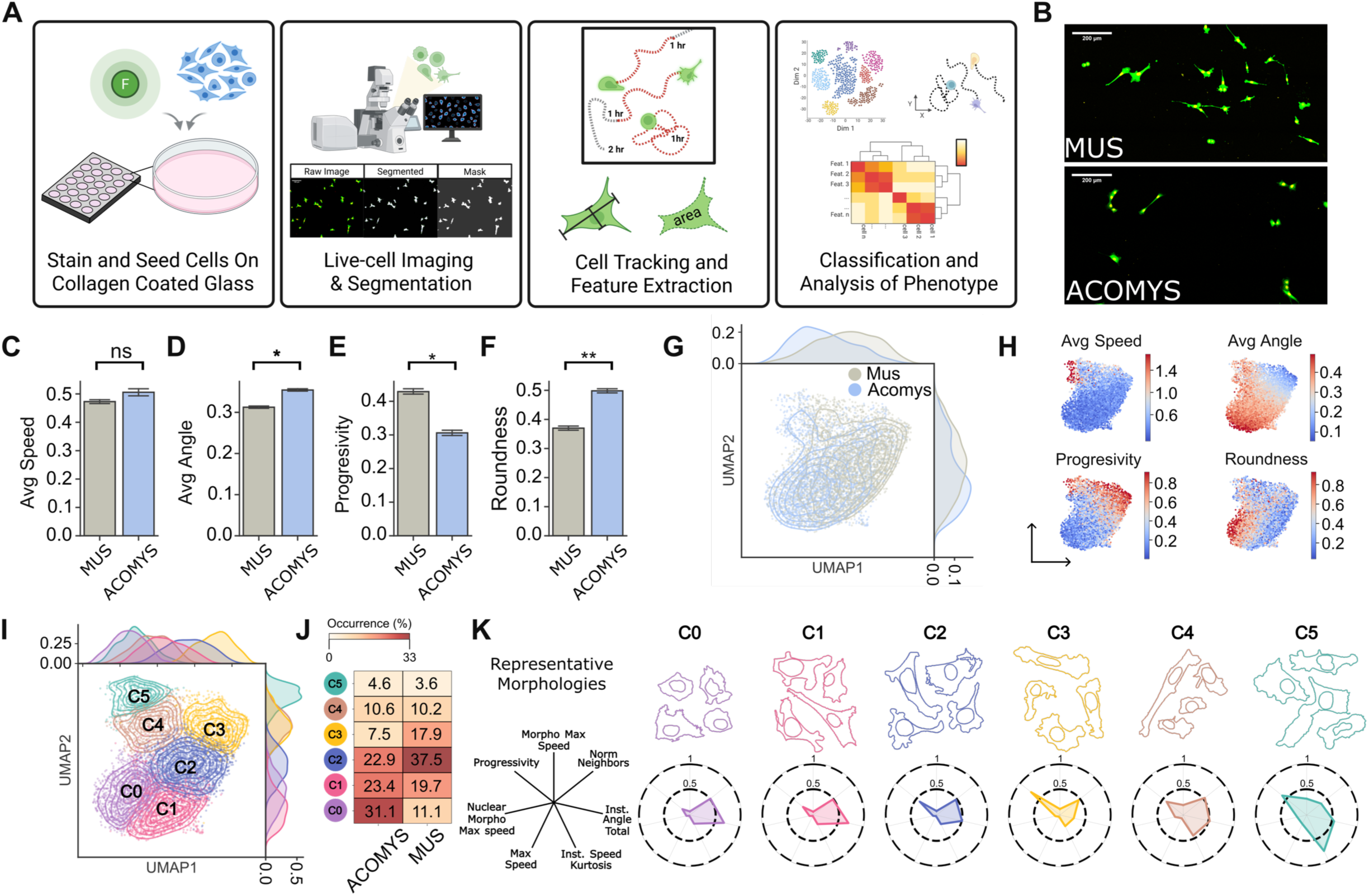
Morphodynamic behavior of *Acomys* and *Mus*. (A) Schematic overview of the experimental and computational pipeline. Cells were stained and seeded onto collagen-coated glass and then underwent live-cell imaging and image segmentation. Single-cell tracking and feature extraction were used to quantify dynamic cell behaviors and morphology. The resulting feature set was used for phenotype classification and analysis. Created with BioRender.com (B) Representative segmented live-cell images show distinct cellular phenotypes for *Acomys* and *Mus*. (C) Bar graphs illustrate average speed (μm/min), average angle (radians), roundness, and progressivity. (D) UMAP space shows the single-cell trajectory behavior (left) and the 2D morphology features: roundness, average speed, average angle, and progressivity as they interface with the UMAP (right). (E) UMAP projection of single-cell phenotypic states identifying six K-means clusters, with accompanying cluster occurrence frequencies in *Acomys* and *Mus*. (F) Representative radar plots encapsulating the morphological feature profiles for each phenotypic cluster and their top four representative morphologies. Effect sizes are reported as Cohen’s d: *small, d ≥ 0.2; **medium, d ≥ 0.5; ***large, d ≥ 0.8. “ns” indicates effect size below 0.2. N=3, n=6228.

The two species displayed comparable migration speeds in 2D (**Figure 1C**). However, *Acomys* cells exhibited a wider range of turning angles (**Figure 1D**) which is reflected in reduced directional progressivity (**Figure 1E**). *Mus* cells displayed more elongated, triangular, and ruffled-edge morphologies compared with the more rounded appearance of *Acomys* cells (**Figure 1F**). *Mus* cells had average cellular diameters of 129.65 µm, whereas *Acomys* cells averaged 140.35 µm. To compare the complete biophysical phenotypes, we first performed Principal Component Analysis (PCA) on more than 90 extracted features describing cell and nuclear morphology and motility, which explained 95% of the total variance (Greenacre et al., 2022). The resulting principal components were used for Uniform Manifold Approximation and Projection (UMAP) visualization, where each dot represents a cell-nucleus pair (**Figure 1G**). Despite both fibroblast lines being of murine origin, UMAP coordinates separated by biophysical phenotypes and species. The upper right of the manifold contained predominantly *Mus* data points, whereas the lower left consisted almost exclusively of *Acomys* cells. The features contributing to this separation included average speed, cell roundness, average turning angle, and progressivity (**Figure 1H**). A UMAP representation of select features are shown in **Supplemental Figure 1**. Together, these results indicate that *Acomys* fibroblasts occupy a distinct biophysical state characterized by reduced directional persistence, increased morphological variability, and unique motility patterns relative to *Mus* fibroblasts.

To characterize the cellular heterogeneity and distribution patterns underlying the biophysical behaviors of the two species, we identified subpopulations of cells sharing similar phenotypes using k-means clustering (**Figure 1I**). These clusters also revealed species-specific phenotypes. Clusters C0 and C1 were largely composed of *Acomys* fibroblasts, whereas cluster C2 was predominantly populated by *Mus* cells (**Figure 1J**). Cluster 0 exhibits a significantly reduced cytoplasmic area, resulting in a higher nuclear-to-cytoplasmic ratio and tightly clustered cell boundaries. Although the cluster labels were ordered by speed and *Mus* cells appear to dominate the later clusters, the fastest migratory cluster (C5) contains *Acomys* cells that migrate faster than the *Mus* cells within the same cluster. The outliers of cell speed can be seen when plotting all individual cells from both species (**Supplemental Figure S2**). The greater proportion of stationary *Acomys* cells is consistent with the bulk properties shown in Figure 1C. Because each k-means cluster represented a unique morphodynamic phenotype, we extracted representative morphologies for each cluster and generated a biophysical fingerprint based on a subset of features, which we represented as a radar plot (**Figure 1K**). Each cluster showed a distinct radar profile.

### Col3A1 perturbation shows limited impact on 2D biophysical phenotypes in *Acomys* fibroblasts

Given the established role of type-III collagen in ECM organization and wound healing, its observed upregulation in *Acomys*, and its implications for Col1 assembly (Allen & Seifert, 2025), we sought to determine whether the gene encoding type-III collagen, Col3A1, contributes to the distinct biophysical phenotypes of *Acomys* fibroblasts. We generated a CRISPR/Cas9-mediated Col3A1 knockout (KO) line using lentiviral third-generation vectors encoding SpCas9 and guide RNAs targeting exon 1 and exon 2 (**Figure 2A**). Successful knockout was confirmed through antibiotic selection, enabling direct comparison between wild-type (WT) and Col3A1-deficient cells.

**Figure 2.**
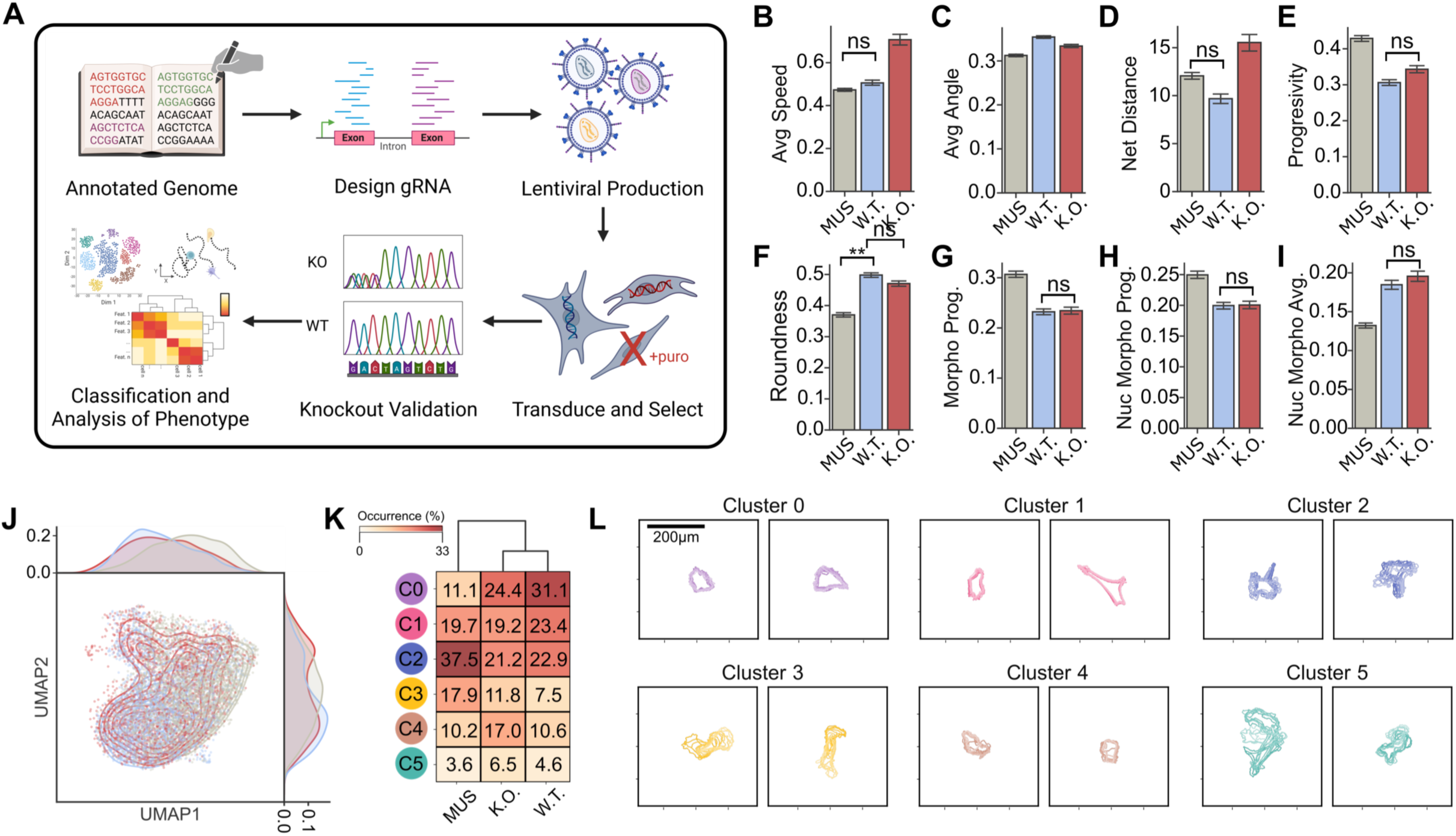
Knockout morphodynamic behavior of *Acomys* and *Mus*. (A) Schematic of CRISPR-based knockout workflow. Annotated phenotypic analyses informed gRNA design, followed by lentiviral production, cell transduction and puromycin selection, knockout validation, and phenotypic classification. Created BioRender.com (B-I) Quantification of cell motility and morphology in *Mus*, *Acomys* wild-type (WT) and knockout (KO) cells. Bar plots show average speed (μm/min), average angle (radians), roundness, progressivity, net distance, morphological progression, nuclear morphological progression, and nuclear morphological average. \*\**D>=0.5*; ns, not significant. (J) UMAP representation of WT and KO cells reveals shifts in the overall phenotypic distribution. (K) Heatmap shows cluster occurrence frequencies in Mus, *Acomys* WT, and *Acomys* KO cells. (L) Representative cell morphomotion from selected clusters illustrates characteristic morphological states for all cell lines, units are pixels. N=3, n=8475.

We then evaluated the impact of Col3A1 loss on 2D cellular motility and morphology using the biophysical assay framework described above. Surprisingly, Col3A1 knockout produced minimal changes in morphological features such as: average turning angle, roundness, and protrusive activity. However, Col3A cells exhibited distinct motility characteristics. KO cells showed reduced substrate adhesion, leading to increased detachment and rolling behavior. This shift in adhesion dynamics resulted in altered displacement patterns and increased apparent migration distances (**Figure 2B**). To compare dynamic morphological profiles, we projected all morphological features into principal component space and quantified single-cell translocation through this space over time. In addition to motility-related features (**Figure 2B–E**), this analysis captures variation in average roundness and motility within the principal component space (**Figure 2F–I**).

To further contextualize these findings, we projected *Acomys* WT, KO, and *Mus* fibroblasts into the UMAP space developed previously (**Figure 2J**). *Acomys* WT and KO populations occupied overlapping regions of the manifold, maintaining clear separation from *Mus* fibroblasts. This finding indicates that loss of Col3A1 does not significantly alter the biophysical identity of *Acomys* fibroblasts. Likewise, clustering analysis showed that WT and KO *Acomys* cells grouped similarly, while *Mus* cells preferentially occupied distinct clusters as before (**Figure 2K**). To investigate behavioral differences, we analyzed representative cell trajectories within clustered subpopulations. Cells in cluster C2, dominated by *Mus* fibroblasts, exhibited highly polarized, fan-shaped morphologies with coordinated wave-like protrusions, indicative of directional cytoskeletal remodeling. In contrast, cluster C0 cells, primarily *Acomys*, displayed limited motility and more isotropic, non-polarized shape fluctuations (**Figure 2L**). These patterns remained consistent across both WT and Col3A1 KO *Acomys* populations. Unsurprisingly, these results indicate that while *Col3A1* likely influences adhesion-related motility behaviors, it does not drive the primary biophysical distinctions between *Acomys* and *Mus* fibroblasts. Instead, the characteristic phenotype of *Acomys* cells appears to be modulated by broader cellular properties that are largely independent of *Col3A1* expression in 2D environments.

### *Acomys* fibroblasts display resistance to senescence induction with unique morphological and proteomic signatures

Previous attempts to model senescence in *Acomys* fibroblasts have been unsuccessful due to poor induction efficacy. Saxena et al. (Saxena et al., 2019) attempted to induce senescence via oxidative stress with H_2_O_2_ without success. Suda et al. (Suda et al., 2024) damaged the plasma membrane with sodium dodecyl sulfate, resulting in extensive cell apoptosis, but the authors identified no concentration that consistently induced senescence. Given the scarless healing observed in *Acomys*, we investigated whether *Acomys* fibroblasts exhibit distinct responses to senescence-inducing stress.

We first evaluated multiple induction strategies. Consistent with prior reports, oxidative stress via H_2_O_2_ failed to reliably induce stable senescent phenotypes in *Acomys* fibroblasts. We then tested bleomycin at concentrations of 250, 200, 100, 50, and 10 μg/mL. However, senescence induction was weak and inconsistent. Finally, we induced DNA damage with the chemotherapy drug doxorubicin. Cells were exposed to doxorubicin for 24 hours and then cultured under normal conditions for 7 days before fixation and imaging (**Figure 3A-B**). Representative fixed and stained cells from day 7 are shown in Figure 3B. *Acomys* and *Mus* doubling rates were confirmed under normal culture conditions (Figure 3C). After DOX addition, *Acomys* underwent a complete senescent transition from DNA damage as measured from cessation in proliferation, however, this required a significantly higher drug concentration compared with *Mus*. Quantification of proliferation rates over 7 days showed that *Acomys* required a 5-fold higher concentration of doxorubicin than *Mus* to achieve a complete halt of proliferation (**Figure 3D**). Other attempts with bleomycin achieved comparable halts in proliferation, but escaper cells resulted in incomplete induction after 7 days of culture.

**Figure 3.**
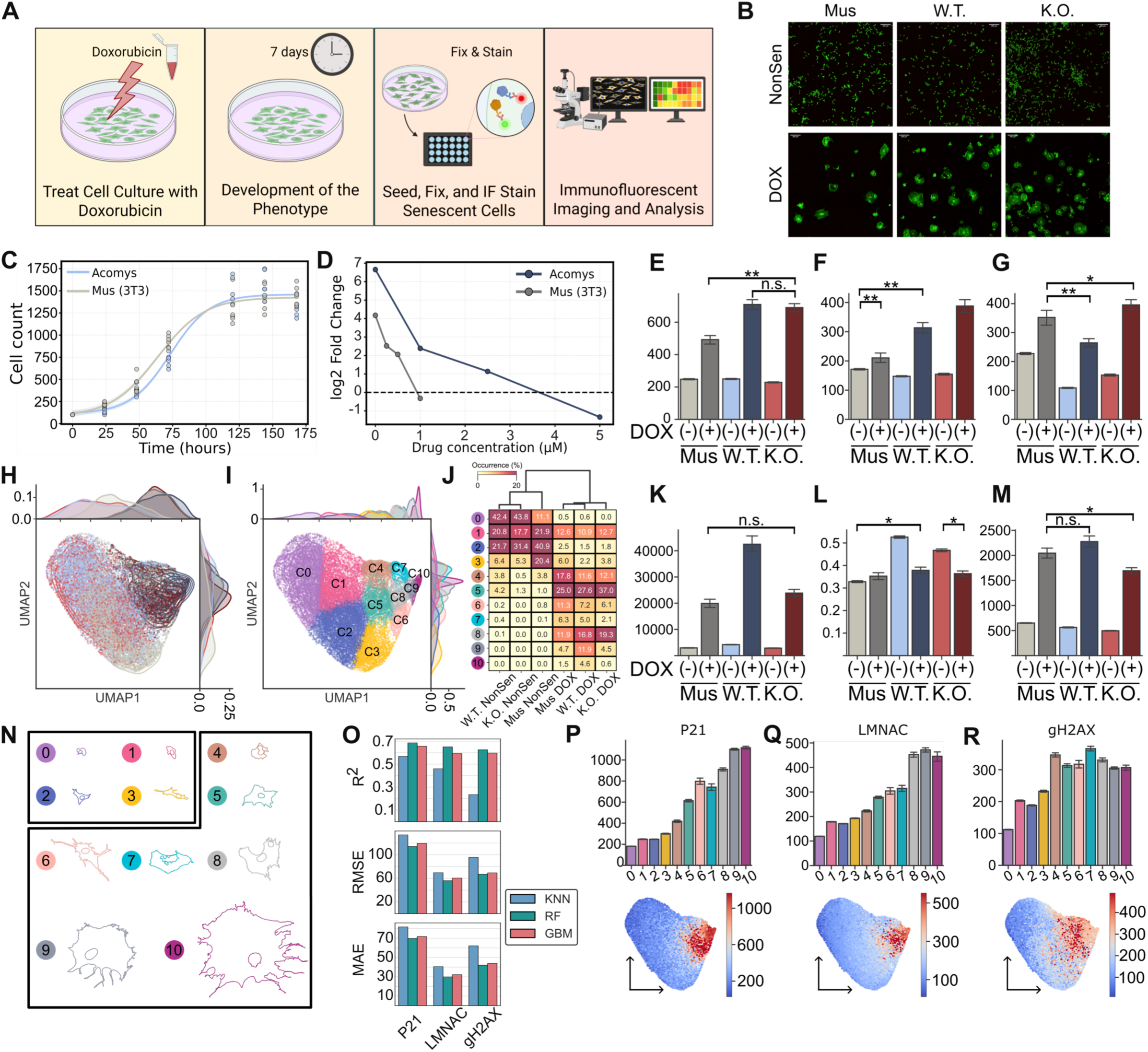
Doxorubicin-induced senescence shapes cellular phenotype and demonstrates species- and genotype-dependent morphological states. (A) Experimental workflow for induction and analysis of senescence. Cells were exposed to doxorubicin (DOX), cultured for 7 days to allow phenotype development, then seeded, fixed, stained, and analyzed by fluorescent imaging. (B) Immunofluorescence images of *Mus*, wild-type (WT), and knockout (KO) cells under control and DOX-treated conditions show condition-dependent changes in cell morphology and senescence-associated features. (C) Cell growth curves for *Acomys* and *Mus* cultures. (D) Log2 fold-changes in cell number across increasing DOX concentrations show differential drug sensitivity between species. (E) Single-cell measurements of cellular p21, nuclear LMNAC, and nuclear γH2AX across untreated and DOX-treated conditions in *Mus*, *Acomys* WT, and KO cells. (F) UMAP space showing the global distribution of untreated and DOX-treated cells. (G) Phenotypic clustering identifies discrete cell states; the heatmap summarizes cluster occurrence frequencies across experimental conditions. (H) Quantification of cell area, cell circularity, and nuclear area across *Mus*, WT, and KO cells with and without DOX treatment. (I) Representative morphologies illustrate the characteristic shapes associated with the identified phenotypic clusters. (J) Bar graph represents comparison of model performance metrics for prediction of p21, LMNAC, and γH2AX levels from morphological features. (K) Top panel represents mean p21, LMNAC, and γH2AX signal across phenotypic clusters. Bottom panel represents UMAP feature maps that show the spatial distribution of these marker intensities across the single-cell state space. IF, immunofluorescence. N=4, n=20622.

To confirm senescent induction, we assessed established morphological and molecular markers (Kudlova et al., 2022). After doxorubicin treatment, both *Acomys* and *Mus* fibroblasts increased in size and morphological heterogeneity, as expected. To quantify biomarker expression across conditions, we batch-corrected fluorescence intensities for single cells to account for variability in imaging and differences in cell and nuclear size. Consistent with senescence, p21 expression increased in both species. Although total Lamin A/C intensity appeared elevated, area-normalized analysis revealed a decrease in Lamin A/C expression; γH2AX levels also increased after DNA damage. *Acomys* fibroblasts exhibited significantly lower γH2AX levels than *Mus* fibroblasts (**Figure 3E-G**), suggesting reduced accumulation or more efficient resolution of DNA damage, even at elevated drug concentrations. Together, these results demonstrate that *Acomys* fibroblasts undergo senescence but require stronger induction and exhibit a distinct damage-response profile. This elevated resistance and altered DNA damage signaling may represent key cellular features underlying regenerative, rather than fibrotic, tissue outcomes.

Building on previous work (Kamat et al., 2024), we quantified the biophysical properties and biomarker expression profiles of senescent cells to determine whether they could serve as a classification and predictive tool. Given the limited temporal resolution of fixed-cell imaging, we employed the high-throughput cell phenotyping pipeline for extracting an expanded cell and nuclei feature set (Wu et al., 2015). Notably, cellular and nuclear sizes increased, and shapes became more heterogeneous. As in Figures 1 and 2, we used principal components as input for UMAP visualization and K-means clustering (**Figure 3H-I**). While morphology shifted between species, a more pronounced shift occurred upon senescence induction. Senescent cells shifted toward the right of the manifold, reflecting larger, more extreme morphologies. Of the 11 k-means clusters, uninduced populations were underrepresented, and most unique morphologies reflected the extreme heterogeneity of doxorubicin-treated cells (**Figure 3J**). Major variance drivers for each species before and after induction are shown in **Figure 3K-M**. Representative morphologies from each cluster revealed non-induced cells in clusters C0, C1, C2, and C3 and primarily doxorubicin-induced cells in clusters C4 through C10 (**Figure 3N**). Strikingly, cluster C4 consisted of cells that seemed to have been arrested in division upon receiving sufficient DNA damage to induce senescence; the unique morphology of a dividing nucleus was preserved.

### Senescent populations for both *Acomys* and *Mus* show discrete subtypes

After staining cells exposed to doxorubicin, we noticed that certain single-cell variations lack one or more positive biomarkers despite similar levels of DNA-damaging agents. Curious whether the morphology-defined space alone could maximize the classification among subtypes, we investigated some conventional senescence biomarkers (p21, LMNAC, and γH2AX) alongside morphological profiles (Kudlova et al., 2022). Because microscopy has a constraint of four channels for imaging, we were limited to two channels for biomarker measurements alongside Hoescht and phalloidin for nuclear and actin structures, respectively. Next, we applied a morphology-based biomarker imputation framework and evaluated three complementary regression architectures that infer molecular expressions directly from single-cell morphological features (**Figure 3O**).

K-Nearest Neighbors predicts biomarker intensity by identifying phenotypically similar cells in a high-dimensional morphological feature space and computing a distance-weighted average of their known biomarker values (Altman, 1992). This nonparametric approach preserves local structure but is sensitive to feature scaling and neighborhood density. To address these limitations, we also employed Gradient Boosting Machine and Random Forest models. All models were trained using morphology alone as input features and benchmarked on the same held-out cells with known biomarker values. Model performance was assessed using R², RMSE, and MAE metrics on the test set (**Figure 3O**).

With both morphological parameters and biomarker expression profiles for each cell, we sought to identify distinct fibroblast subtypes. Strikingly, of the senescent morphological subtypes, clusters C4 and C7 exhibited significantly higher total γH2AX expression than even the largest cells (**Figure 3P-R**). These results indicate that senescence-associated morphology and DNA damage signaling are not linearly coupled but rather define functionally distinct cellular states.

Morphological features such as cell area, roundness, and protrusive structure primarily distinguished enlarged, structurally remodeled senescent cells, whereas γH2AX intensity distinguished cells with elevated DNA damage independent of size. Thus, the population contains at least two separable senescent subtypes: a morphology-dominant state that is positive for traditional senescence markers and correlates with cell size, and a damage-associated, size-independent state. Each subtype is defined by distinct feature contributions. This separation suggests that senescence is not a uniform phenotype but rather a spectrum of states that may differ in functional behavior and response to perturbations.

### Enhanced matrix remodeling in *Acomys* is promoted by SASP-driven senescent cells

To investigate fibroblast behavior in a wound-remodeling context, we assessed *Acomys* and *Mus* cells using 2D scratch assays and 3D collagen gel contraction assays. The 2D scratch wound closure assay enabled continuous evaluation of cell migration via monolayer closure over 12 hours (**Figure 4A**). **Figure 4B** shows representative images of progressive wound closure at 0, 200, 400, and 600 minutes post-scratch for *Acomys* cells. Compared with *Mus* fibroblasts, *Acomys* WT fibroblasts did not differ significantly in scratch closure under standard culture conditions (**Figure 4C**). Although the 2D scratch assay provided an initial model for migratory behavior, we extended this analysis to a 3D context to more directly measure physiological remodeling capacity and to decouple proliferation from migration.

**Figure 4.**
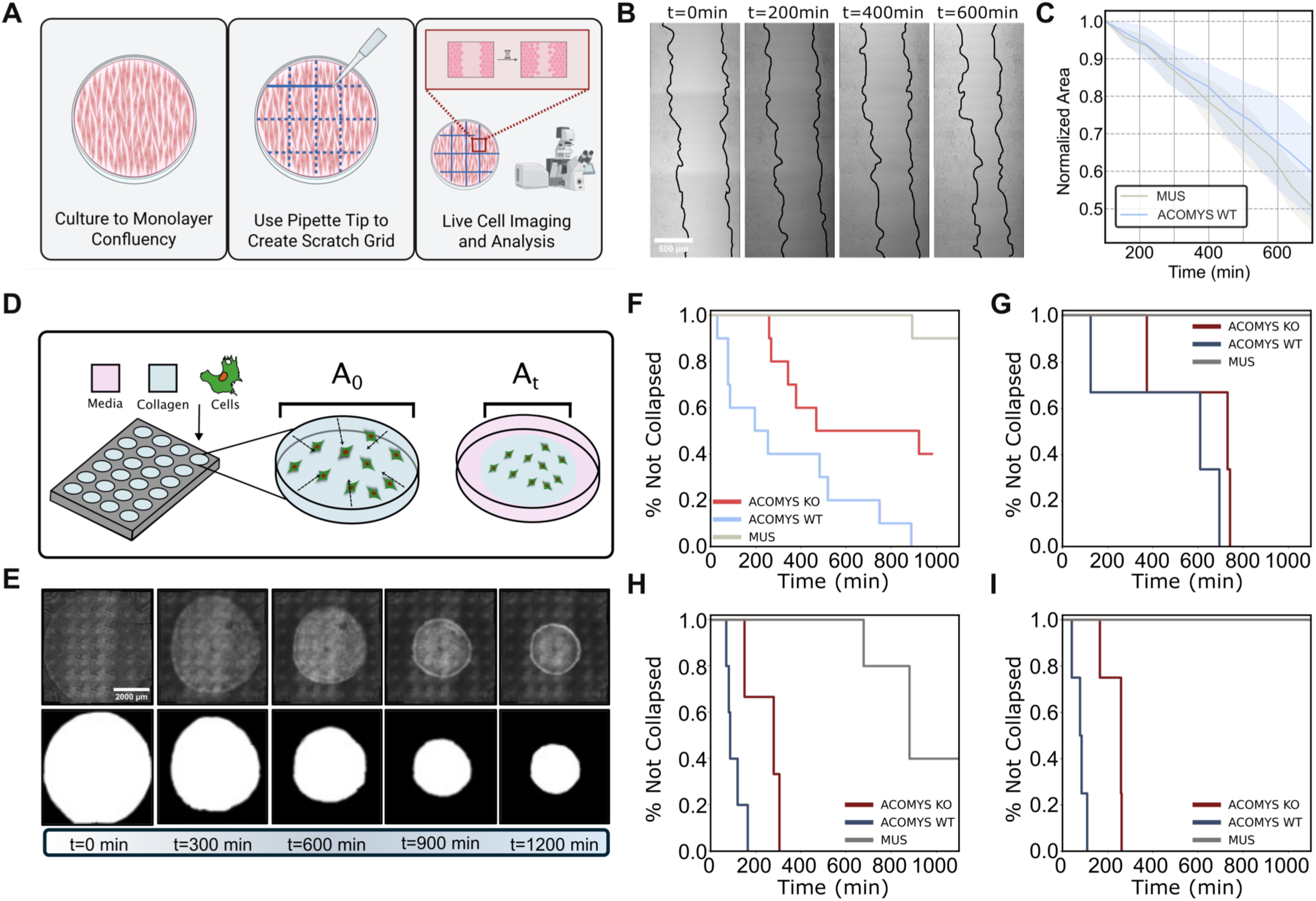
Comparative Analysis of Tissue Remodeling Capacity of *Acomys* and *Mus* and the Effects of Senescent Cells and SASP on Collagen Gel Collapse Dynamics. (A) Schematic of the live-cell scratch assay. Confluent monolayers were mechanically wounded with a pipette tip, and wound closure was quantified by time-lapse imaging. Made with BioRender.com (B) Time-lapse images show progressive wound closure at indicated timepoints during cell migration. (C) Normalized wound areas over time for *Mus* and *Acomys* wild-type cells show differences in migration dynamics. (D) Schematic of the collagen gel collapse assay in which cells were embedded in collagen matrices and gel collapse was monitored over time. (E) Representative images of collagen gel collapse. Time-course brightfield images and corresponding segmented masks show progressive reduction in gel area during matrix contraction. (F-I) Kaplan-Meier plots show the fraction of gels remaining intact over time across non-senescent *Acomys* and *Mus* conditions (F), doxorubicin-exposed conditions (G), non-senescent conditions with 10% doxorubicin (H), and non-senescent cells with conditioned media (I), revealing condition- and cell-dependent differences in collagen gel collapse kinetics.

We established a collagen gel contraction assay as a proxy for matrix remodeling and contractile force generation. Cells were embedded within a collagen matrix that polymerized to fill the entire well area. Gel contraction was quantified over time by measuring area reduction as cell-generated traction forces induced gel compaction and detachment from the well perimeter (**Figure 4D-E**). We hypothesized that *Acomys* fibroblasts would remodel faster than *Mus*, resulting in more rapid gel contraction. Brightfield images of the entire well were captured to visualize and segment the collagen gel area, and the area of each frame was normalized to the initial frame. *Acomys* WT cells exhibited a marked increase in contraction speed. Plotting the time to reach two-thirds of the original gel area showed that *Mus* cells took 900 to 1200 minutes, whereas *Acomys* WT cells required just 20 to 900 minutes across replicates. Interestingly, the Col3A1 KO line exhibited reduced remodeling potential, contracting at a rate between that of *Mus* and *Acomys* WT (**Figure 4F**).

Next, we examined whether senescent cells exhibited altered contraction dynamics compared with non-induced populations. Using doxorubicin-induced cells alone, contraction speed was comparable to non-induced controls (**Figure 4G**). To mimic acute wound conditions, we tested a 10% senescent population mixed with non-induced cells for each species. Surprisingly, contraction speed increased further (**Figure 4H**). This finding suggested that cell-cell interactions may modulate matrix remodeling or that secreted factors influence baseline cell behavior. To test these possibilities, we collected conditioned media from 24 hours of culture with 100,000 7-day doxorubicin-induced cells. We then applied diluted conditioned media to baseline fibroblasts and observed an acceleration of gel contraction similar to that of the co-culture (**Figure 4I**). This finding suggests that SASP factors may contribute to enhanced remodeling capacity.

## DISCUSSION

By characterizing an immortalized *Acomys* fibroblast line and its Col3A1 KO derivative, we developed a platform to investigate cellular mechanisms that may contribute to regenerative wound healing. Our comparative analyses revealed that *Acomys* WT fibroblasts exhibit distinct morphological features, including increased cell diameters and reduced complexity compared with *Mus* fibroblasts, while their motility remained comparable. These properties may contribute to the regenerative capabilities of *Acomys*, consistent with previous reports of distinct fibroblast phenotypes in this species.

Our findings demonstrate that fibroblast state, defined by biophysical properties, ECM interactions, and senescence-associated signaling, plays a central role in shaping matrix remodeling outcomes. The dissociation between morphology, DNA damage signaling, and functional behavior suggests that senescence is not a uniform endpoint but rather a spectrum of states with distinct consequences. Similarly, the context-dependent effect of Col3A1 loss indicates that matrix remodeling capacity arises from integrated cellular programs rather than any single ECM component. This study positions *Acomys* fibroblasts as a tractable model for studying fibroblast states that differ from fibrotic responses in *Mus*. *Acomys* cells have been reported to resist H_2_O_2_-induced senescence in part by limiting oxidative stress and ROS-associated damage, yet here we show they remain capable of mounting a senescence response to direct genotoxic injury from doxorubicin. This suggests that senescence induction in *Acomys* may be stimulus dependent rather than globally suppressed. Together with the presence of distinct senescent subtypes, these features warrant further investigation into how cellular heterogeneity and DNA damage responses shape fibrotic and tissue-level outcomes. These features may offer advantages over existing mouse models for studying fibroblast behavior where regeneration and fibrosis diverge.

Although our findings were derived from *ex vivo* systems, they provide a foundation for understanding how fibroblast states may contribute to tissue dynamics in vivo. The modulation of collagen matrix contraction by senescent cell-derived factors suggests that paracrine signaling within heterogeneous cell populations could influence matrix organization during wound healing. Future studies integrating in vivo models will be essential to determine how these cellular states translate to physiological repair. Our results define a distinct *Acomys* fibroblast phenotype characterized by altered biophysical behavior, resistance to senescence induction, and enhanced matrix remodeling. These properties establish a platform for investigating non-fibrotic fibroblast states and developing strategies to promote regenerative outcomes in mammalian tissue repair.

## RESOURCE AVAILABILITY

All data supporting the findings in this study are presented in the manuscript and its Supplementary documents.

## Data availability

All data supporting the findings of the manuscript are presented in the figures and supplementary materials

## ACKNOWLEDGMENTS

We would like to thank members of the Jain Lab at the University of Florida and members of the Phillip lab for their feedback and support throughout the project. We would like to thank Dr. Michelle Dill for access to immortalized *Acomys* lines, Dr. Malcom Maden and Dr. Chelsey Simmons for securing funding and ideation around repair mechanisms, and Dr. Barbazuk for genome access around Col3A1 design. We acknowledge financial support from the NIH NIGMS MIRA 5R35GM147788 (PKJ), NIH OD 5R21OD028211 (PKJ), NIH NIGMS MIRA R35-GM157099 (JMP), the Salisbury Family and Center for Innovative Medicine Human Aging Project Scholar Award (JMP), and a Catalyst Award from the Johns Hopkins University (JMP).

## AUTHOR CONTRIBUTIONS

Conceptualization: N.M. and J.M.P.; investigation: N.M., M.B., A.L., Y.C., L.N., P.J., and J.M.P.; methodology: N.M., P.J., and J.M.P.; resources: N.M. and J.M.P.; validation: N.M., M.B., A.L., Y.C., and J.M.P.; visualization: N.M. and J.M.P.; data curation: N.M., M.B., A.L., Y.C., and J.M.P.; formal analysis: N.M. and Y.C.; software: N.M. and J.M.P.; supervision: P.J. and J.M.P.; project administration: J.M.P.; writing—original draft: N.M., M.B., A.L., Y.C., and J.M.P.; writing—review and editing: N.M. and J.M.P.; funding acquisition: J.M.P.

## MATERIALS AND METHODS

### Cell culture

Immortalized *Acomys* and *Mus* mouse dermal fibroblasts were procured from the University of Florida [MB6.1] and Johns Hopkins University, respectively, and used as the experimental standard for this study. Three cell lines were used: *Acomys* WT, *Acomys* KO (a Col3A1 collagen KO within *Acomys* species), and *Mus* 3T3. Cells were initially maintained in Dulbecco’s Modified Eagle Medium (DMEM) supplemented with 15% fetal bovine serum (FBS) and 1% penicillin-streptomycin in a humidified atmosphere of 5% CO_2_ at 37°C. Cells were seeded onto cell culture-treated culture dishes or onto collagen-I–coated glass bottom dishes with DMEM at a density of 75,000 cells/cm^2^. After cells were cultured, they were allowed to adhere overnight before being passaged to a new flask. Cells underwent one passage after being thawed from the cryo-freezer with passage numbers ranging from 30 to 35 for 3T3 *Mus* cells and 6 to 15 for *Acomys* WT and KO cells.

### Live-cell imaging and microscopy

To assess the physical behavior of these cells, we first seeded low-density fibroblasts from each species onto collagen-coated glass plates and stained the nuclear DNA and actin with live-cell dyes according to manufacturer protocols (SPY555-DNA Cat. #CY-SC201 for nuclear and SPY650-FastAct Cat. #CY-SC505 for actin). We then used confocal microscopy to observe single-cell shapes and positions every 5 minutes for at least 4 hours.

Cells were imaged on a confocal fluorescence microscope (Leica Stellaris 5) equipped with a live-cell imaging chamber (Tokai Hit), which maintained optimal culturing conditions at 37°C with 5% CO_2_ when necessary. For live-cell movies, we captured fluorescence images at regular intervals (every 5 minutes) over the course of ∼4 hours to visualize cell and nuclear morphology. Live-cell images were captured using a 20X objective with 2X optical zoom. Fixed-cell images were also acquired with the Leica Stellaris 5, and fluorescent images were taken at 20X magnification with four different laser lines (405 Diode, 488 Diode, 568 Diode, and 647 Diode). All images were captured at a 1024 x 1024-pixel resolution with 0.568 microns per pixel for 20X, or 0.284 microns per pixel for 40X.

### Segmentation and in-house computational analyses

The acquired images were subjected to CellProfiler segmentation (Stirling et al., 2021). Subsequent instance-segmented masks were then fed to an in-house post-processing pipeline to delineate individual cells (removing all cells touching three or more neighboring cells) and potentially extract more relevant morphological features using pipelines as in a previous study (Phillip et al., 2021). By doing so, we ensured single-cell quantification throughout the captured data. Subsequent data processing techniques are outlined below.

### UMAP and K-means clustering

Across all biological conditions and replicates for the *Acomys* and 3T3 *Mus* cell lines, a reduced dimensionality framework was constructed with the UMAP technique. All morphological construction parameters were standard-scaled after being log-normalized and were subsequently fed into PCA to conserve 95% variance. These generated PCs were used to create a 2D UMAP space. UMAP, a nonlinear dimensionality reduction technique, captures and projects the high-dimensional data structure within a lower-dimensional space for comprehensibility. Within the UMAP space, each cell is represented as a distinct point. In addition to the morphological analysis performed with the UMAP technique, we used K-means clustering, a machine learning algorithm that groups cells based on shared characteristics, to analyze cell morphologies. K-centroids are first initialized randomly within the data space. The model then assigns each data point to the nearest centroid, forming K-clusters. After assignment, the centroids are recalculated as the mean of all points in their respective clusters. This process of assigning points and recalculating centroids repeats iteratively until the centroids stabilize, indicating that the cells either no longer move significantly or a maximum number of iterations has been reached.

### Generation of a Col3A1 KO AcoSI-1 cell line

CRISPR-Cas9 guide RNA targeting Col3A1 was designed with CRISPick (Broad Institute) (Doench et al., 2016). Four guide RNAs were selected (two guides targeting exon 1 and two guides targeting exon 2) and cloned into lentiCRISPR v2 backbone (Addgene plasmid #52961) (Sanjana et al., 2014). For lentiviral packaging, we co-transfected 10 µg of lenCRISPR v2 containing guide RNAs into 5 µg of pMD2.G (pMD2.G was a gift from Didier Trono [Addgene plasmid # 12259; http://n2t.net/addgene:12259; RRID:Addgene_12259]) and 7.5 µg of psPAX2 (psPAX2 was a gift from Didier Trono [Addgene plasmid # 12260; http://n2t.net/addgene:12260; RRID:Addgene_12260]) in a T75 flask using lipofectamine 3000 (Invitrogen, Cat# L3000001) according to the manufacturer’s protocol. Six hours later, medium was replaced with lentiviral harvesting medium (DMEM, 10% FBS, 1x penstrep, and 1% bovine serum albumin [BSA]) and incubated for 60 hours at 37°C and 5% CO_2_. Next, the medium was collected, centrifuged, and passed through a 0.45 µm filter.

Lentiviral particles were introduced into AcoSI-1 cells by reverse transduction at a multiplicity of infection of 0.5 in 10 µg/mL polybrene. Cells were incubated for 72 hours and then underwent selection with 10 µg/mL puromycin for 7 days. For Col3A1 KO validation, the cells were lysed and analyzed by targeted amplicon sequencing.

### Senescence induction

Drugs used for senescence induction were diluted from stock solutions into DMSO (<0.2% vol/vol) in accordance with manufacturer guidelines. The resulting solutions were subsequently diluted in media to a final working concentration for cell line exposure. We initially tested bleomycin in cell lines for 4 hours at concentrations of 10, 50, 100, 200, and 250 μg/mL but ultimately chose to use the chemotherapeutic drug doxorubicin. Thus, senescence was induced by treating cells for 24 hours with doxorubicin at concentrations of 1 μM for 3T3 *Mus* cells and 5 μM for *Acomys* cells.

To induce senescence, we removed the initial cell culture medium from the dishes and replaced it with induction medium (cell culture medium with respectively proportioned volumes of doxorubicin per cell line). Then, the cells were incubated for 24 hours under the same incubation conditions provided previously. After incubation, the cells were washed with 1X phosphate-buffered saline (PBS; Corning, 21–031) before being supplemented with 8 to 10 mL of fresh cell culture medium.

Cells were allowed to proliferate for 5 to 7 days to allow for development of senescent phenotypes. Next, the cells were plated onto 50 μg/mL collagen I–coated 24-well glass bottom plates at a density of 2000 cells/well (control) and 1250 cells/well (senescent) in 1 mL of cell culture medium. For high-throughput assays in 96-well plates, each cell line was plated at a density of 300 cells/well (control) and 250 cells/well (senescent) in 200 μL of medium per well. After the induction and subsequent development of the senescence phenotype, cells were visualized by live-cell microscopy before being analyzed for additional morphological and functional changes.

### Cellular fixation and immunofluorescence staining

After control and senescent cells were seeded at a density of 2000 cells/well in a 24-well plate, they were allowed to adhere overnight. Once cells had adhered, they were immediately washed with 1X PBS and fixed in freshly prepared 4% (w/v) paraformaldehyde and 1X PBS solution for 10 minutes. After fixation, cells were washed three times with 1X PBS before being permeabilized by incubation in a solution of 0.1% Triton-X in 1X PBS for 10 minutes. Cells were again washed three times with 1X PBS to remove any residual Triton-X solution before being blocked with 2% BSA (weight/volume in 1X PBS) for at least 30 minutes at room temperature. After this blocking step, cells were incubated with primary antibodies, including LMNAC (Abcam, ab224816, 1:500) and γH2AX (Abcam, ab81299, 1:250) in 1% BSA overnight at 4°C. After 16 to 18 hours, cells were washed three times with 1X PBS and incubated in p21 (Santa Cruz, SC-817, 1:200) in 1% BSA for 1.5 hours at room temperature. Cells were washed three times with 1X PBS to remove any residual primary antibodies before being incubated in a cocktail of secondary antibodies and small molecule fluorophores, including Hoechst (Thermo Fisher, H3570, 1:250), Alexa Fluor-488 phalloidin (Thermo Fisher, A12379, 1:200), Alexa Fluor-647 phalloidin (Thermo Fisher, A12379, 1:200), goat anti-rabbit IgG-Alexa Fluor 568 (Thermo Fisher, A-11036, 1:500), goat anti-mouse IgG-Alexa Fluor 488 (Thermo Fisher, A-21240, 1:500), and goat anti-mouse IgG-Alexa Fluor 647 (Thermo Fisher, A-21240, 1:500) in 1% BSA. After incubation in secondary antibodies, the cells were washed three times with 1X PBS and imaged on a Leica Stellaris 5 confocal microscope. All samples were imaged immediately after staining or within 24 hours of secondary antibody incubation, with intermediate storage at 4°C.

### Phenotype imputation

Gradient Boosting Machines model nonlinear relationships by iteratively fitting ensembles of shallow decision trees, where each successive tree corrects the residuals of the previous model (Natekin & Knoll, 2013). This approach is well-suited for capturing complex, nonadditive dependencies between morphology and biomarker expression while maintaining strong generalization performance. Random Forest regression constructs an ensemble of decorrelated decision trees trained on bootstrapped samples of data and random subsets of features (Couronné et al., 2018). By averaging across trees, Random Forest reduces variance and overfitting while modeling high-order interactions between morphological descriptors and molecular state.

### Scratch assay

To evaluate wound healing capacity and collective migration behavior, we performed a 2D scratch assay in *Acomys* WT and *Mus* 3T3 fibroblasts under control conditions. Cells were seeded in 24-well tissue culture plates at a density of 75,000 per well in 1 mL of standard growth medium and grown to full confluence over 24 to 48 hours. Upon reaching confluence, cells were gently washed twice with 1X PBS to remove any nonadherent or dead cells. A uniform scratch was introduced through the center of each well with a 200 µL pipette tip. The plates were washed once with PBS to remove detached cells, and each well was replenished with fresh growth medium.

We imaged plates at regular intervals post-scratch on a phase-contrast inverted microscope. We kept imaging parameters such as magnification, exposure, and position constant to ensure comparability across timepoints and conditions. We analyzed scratch areas using ImageJ by tracing the wound boundaries and calculating the remaining wound area at each timepoint. Percent wound closure was calculated with the following formula:

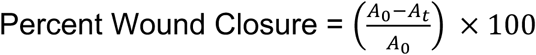

where *A_0_* is the scratch area at time 0 and *A_t_* is the scratch area at the indicated timepoint (24 or 48 hours). We tested each condition in biological triplicates and analyzed results by two-way ANOVA with Tukey’s post hoc test to assess the effect of genotype on wound closure.

### Gel collapse

We performed collagen gel collapse assays to assess matrix remodeling potential in *Acomys* WT, *Acomys* KO, and 3T3 *Mus* cell lines under non-senescent (control) and co-culture conditions. A type I collagen gel master mix was prepared on ice in a sterile 15 mL tube by combining HBSS, DMEM, and NaOH, with pH adjusted to 7.0. Collagen was gradually added with gentle mixing to ensure homogeneous incorporation and preserve structural integrity. Each gel received 55,000 cells per well in 100 µL of master mix. For co-culture wells, we seeded cells at a ratio of 90% non-senescent to 10% senescent fibroblasts to model paracrine interactions within a predominantly healthy population. Plates were incubated for 1 hour at 37°C to allow collagen polymerization and initial cell-matrix interaction, after which each well was hydrated with 100 µL of DMEM. Gels were imaged immediately after hydration to capture the initial state. Magnification, timing, and exposure settings were held constant across all sessions.

To isolate the contribution of secreted soluble factors to gel remodeling, we conducted a conditioned media assay in parallel for each cell line. Cells from each line and condition were incubated in 10 mL of DMEM for 24 hours. Conditioned media was then collected, centrifuged to remove cellular debris, and diluted 1:1 with fresh DMEM. Collagen gels were prepared with the same protocol described above containing only healthy cells and hydrated with 100 µL of conditioned media per well in place of standard DMEM. This control enabled us to assess whether secreted factors alone were sufficient to drive matrix remodeling.

All gels were imaged in brightfield on a Leica Stellaris 5 microscope. Because imaging intervals varied across experimental sessions, we multiplied raw frame indices by the session-specific timestep to yield real time in minutes, ensuring comparability across biorepeats. We delineated gel boundaries by manually segmenting each brightfield frame with ImageJ to produce binary masks. Mask area was extracted computationally by using regionprops from the scikit-image library in Python. Prior to area extraction, we applied a 5×5 morphological closing kernel to each binary mask via OpenCV to bridge any discontinuities in the segmented boundary; the largest connected region was retained for all subsequent measurements. Gel area at each timepoint was normalized to the first-frame area of that gel, yielding an adjusted area initialized at 1.0. Collapse was defined as the first timepoint at which adjusted area fell to ≤67% of the original, with the exact crossing time determined by linear interpolation between the last frame above and first frame below threshold. Gels that never reached this threshold were right-censored at their final observation timepoint. Time-to-collapse was modeled by using Kaplan-Meier survival analysis implemented in the Python lifelines library, with duration measured from the first frame. Pairwise log-rank tests were performed between all condition pairs within each cell line, and *p*-values are reported without correction for multiple comparisons.

### Data Analysis: (Statistics and reproducibility & software)

Quantitative data obtained from image analyses were statistically analyzed with Python software packages. Graphical representations of the data were generated in Python, including UMAP plots, heatmaps, proliferation curves, and violin and box plots to visualize differences in senescence characteristics between cell lines. Effect size was determined by using a Cohen’s D with a significance level set at *small D>=0.2, **medium D>=0.5, and ***large D>=0.8.

**Supplemental Figure 1**. UMAP feature magnitude showing the representative set of features classes that contribute to the UMAP space. Principal Component Analysis (PCA) was applied to more than 90 extracted features describing cellular motility and morphology. Resulting principal components were used as input for UMAP to visualize the phenotypic space, where each dot represents a nuclei and cell pair trajectory.

**Supplemental Figure 2**. Box and violin plots indicate outliers of migratory cells. A-C. Box plots of average, max, and min speeds in um/min, respectively. D-F. Violin plots of average, max, and min speeds in um/min, respectively.

## REFERENCES

1. Brant, J. O., Lopez, M.-C., Baker, H. V., Barbazuk, W. B. & Maden, M. A Comparative Analysis of Gene Expression Profiles during Skin Regeneration in Mus and Acomys. PLOS ONE 10, e0142931 (2015).

2. Phillip, J. M., Han, K.-S., Chen, W.-C., Wirtz, D. & Wu, P.-H. A robust unsupervised machine-learning method to quantify the morphological heterogeneity of cells and nuclei. Nat Protoc 16, 754–774 (2021).

3. Kamal, M., Joanisse, S. & Parise, G. Bleomycin-treated myoblasts undergo p21-associated cellular senescence and have severely impaired differentiation. GeroScience 46, 1843–1859 (2023).

4. Stirling, D. R. et al. CellProfiler 4: improvements in speed, utility and usability. BMC Bioinformatics 22, 433 (2021).

5. Wilkinson, H. N. & Hardman, M. J. Cellular Senescence in Acute and Chronic Wound Repair. Cold Spring Harb Perspect Biol a041221 (2022) doi:10.1101/cshperspect.a041221.

6. Childs, B. G., Durik, M., Baker, D. J. & van Deursen, J. M. Cellular senescence in aging and age-related disease: from mechanisms to therapy. Nat Med 21, 1424–1435 (2015).

7. Kudlova, N., De Sanctis, J. B. & Hajduch, M. Cellular Senescence: Molecular Targets, Biomarkers, and Senolytic Drugs. Int J Mol Sci 23, 4168 (2022).

8. Saxena, S., Vekaria, H., Sullivan, P. G. & Seifert, A. W. Connective tissue fibroblasts from highly regenerative mammals are refractory to ROS-induced cellular senescence. Nat Commun 10, 4400 (2019).

9. Saxena, S., Vekaria, H., Sullivan, P. G. & Seifert, A. W. Connective tissue fibroblasts from highly regenerative mammals are refractory to ROS-induced cellular senescence. Nat Commun 10, 4400 (2019).

10. Sczyrba, A. et al. Critical Assessment of Metagenome Interpretation—a benchmark of metagenomics software. Nat Methods 14, 1063–1071 (2017).

11. Wu, P.-H. et al. Evolution of cellular morpho-phenotypes in cancer metastasis. Sci Rep 5, 18437 (2015).

12. Tracy, L. E., Minasian, R. A. & Caterson, E. J. Extracellular Matrix and Dermal Fibroblast Function in the Healing Wound. Advances in Wound Care 5, 119–136 (2016).

13. Li, B. & Wang, J. H.-C. Fibroblasts and myofibroblasts in wound healing: Force generation and measurement. Journal of Tissue Viability 20, 108–120 (2011).

14. Omar, R., Malfait, F. & Van Agtmael, T. Four decades in the making: Collagen III and mechanisms of vascular Ehlers Danlos Syndrome. Matrix Biology Plus 12, 100090 (2021).

15. Natekin, A. & Knoll, A. Gradient boosting machines, a tutorial. Front. Neurorobot. 7, (2013).

16. Sanjana, N. E., Shalem, O. & Zhang, F. Improved vectors and genome-wide libraries for CRISPR screening. Nat Methods 11, 783–784 (2014).

17. Zhang, Z. Introduction to machine learning: k-nearest neighbors. Ann Transl Med 4, 218 (2016).

18. Jin, X. & Han, J. K-Means Clustering. in Encyclopedia of Machine Learning (eds Sammut, C. & Webb, G. I.) 563–564 (Springer US, Boston, MA, 2011). doi:10.1007/978-0-387-30164-8_425.

19. Doench, J. G. et al. Optimized sgRNA design to maximize activity and minimize off-target effects of CRISPR-Cas9. Nat Biotechnol 34, 184–191 (2016).

20. Doench, J. G. et al. Optimized sgRNA design to maximize activity and minimize off-target effects of CRISPR-Cas9. Nat Biotechnol 34, 184–191 (2016).

21. Suda, K. et al. Plasma membrane damage limits replicative lifespan in yeast and induces premature senescence in human fibroblasts. Nat Aging 4, 319–335 (2024).

22. Greenacre, M. et al. Principal component analysis. Nat Rev Methods Primers 2, 100 (2022).

23. Couronné, R., Probst, P. & Boulesteix, A.-L. Random forest versus logistic regression: a large-scale benchmark experiment. BMC Bioinformatics 19, 270 (2018).

24. Saito, Y., Yamamoto, S. & Chikenji, T. S. Role of cellular senescence in inflammation and regeneration. Inflamm Regener 44, 28 (2024).

25. Cialdai, F., Risaliti, C. & Monici, M. Role of fibroblasts in wound healing and tissue remodeling on Earth and in space. Front. Bioeng. Biotechnol. 10, 958381 (2022).

26. Wilkinson, H. N. & Hardman, M. J. Senescence in Wound Repair: Emerging Strategies to Target Chronic Healing Wounds. Front. Cell Dev. Biol. 8, 773 (2020).

27. Kamat, P. et al. Single-cell morphology encodes functional subtypes of senescence in aging human dermal fibroblasts. Preprint at 10.1101/2024.05.10.593637 (2024).

28. Seifert, A. W. et al. Skin shedding and tissue regeneration in African spiny mice (Acomys). Nature 489, 561–565 (2012).

29. Allen, R. S. & Seifert, A. W. Spiny mice (Acomys) have evolved cellular features to support regenerative healing. Annals of the New York Academy of Sciences 1544, 5–26 (2025).

30. Allen, R. S. & Seifert, A. W. Spiny mice (Acomys) have evolved cellular features to support regenerative healing. Ann N Y Acad Sci 1544, 5–26 (2025).

31. Gaire, J. et al. Spiny mouse (Acomys): an emerging research organism for regenerative medicine with applications beyond the skin. npj Regen Med 6, 1 (2021).

32. Kuivaniemi, H. & Tromp, G. Type III collagen (COL3A1): Gene and protein structure, tissue distribution, and associated diseases. Gene 707, 151–171 (2019).

33. Stewart, D. C. et al. Type III Collagen Regulates Matrix Architecture and Mechanosensing during Wound Healing. Journal of Investigative Dermatology 145, 919–938.e14 (2025).

34. McInnes, L., Healy, J., Saul, N. & Großberger, L. UMAP: Uniform Manifold Approximation and Projection. JOSS 3, 861 (2018).

35. Talbott, H. E., Mascharak, S., Griffin, M., Wan, D. C. & Longaker, M. T. Wound healing, fibroblast heterogeneity, and fibrosis. Cell Stem Cell 29, 1161–1180 (2022).

36. Talbott, H. E., Mascharak, S., Griffin, M., Wan, D. C. & Longaker, M. T. Wound healing, fibroblast heterogeneity, and fibrosis. Cell Stem Cell 29, 1161–1180 (2022).

