## Supplementary figures and images for "Behavioral and Functional Profiling of *Acomys cahirinus* Fibroblasts Reveals Enhanced Matrix Remodeling Capacity"

### Supplementary Figures

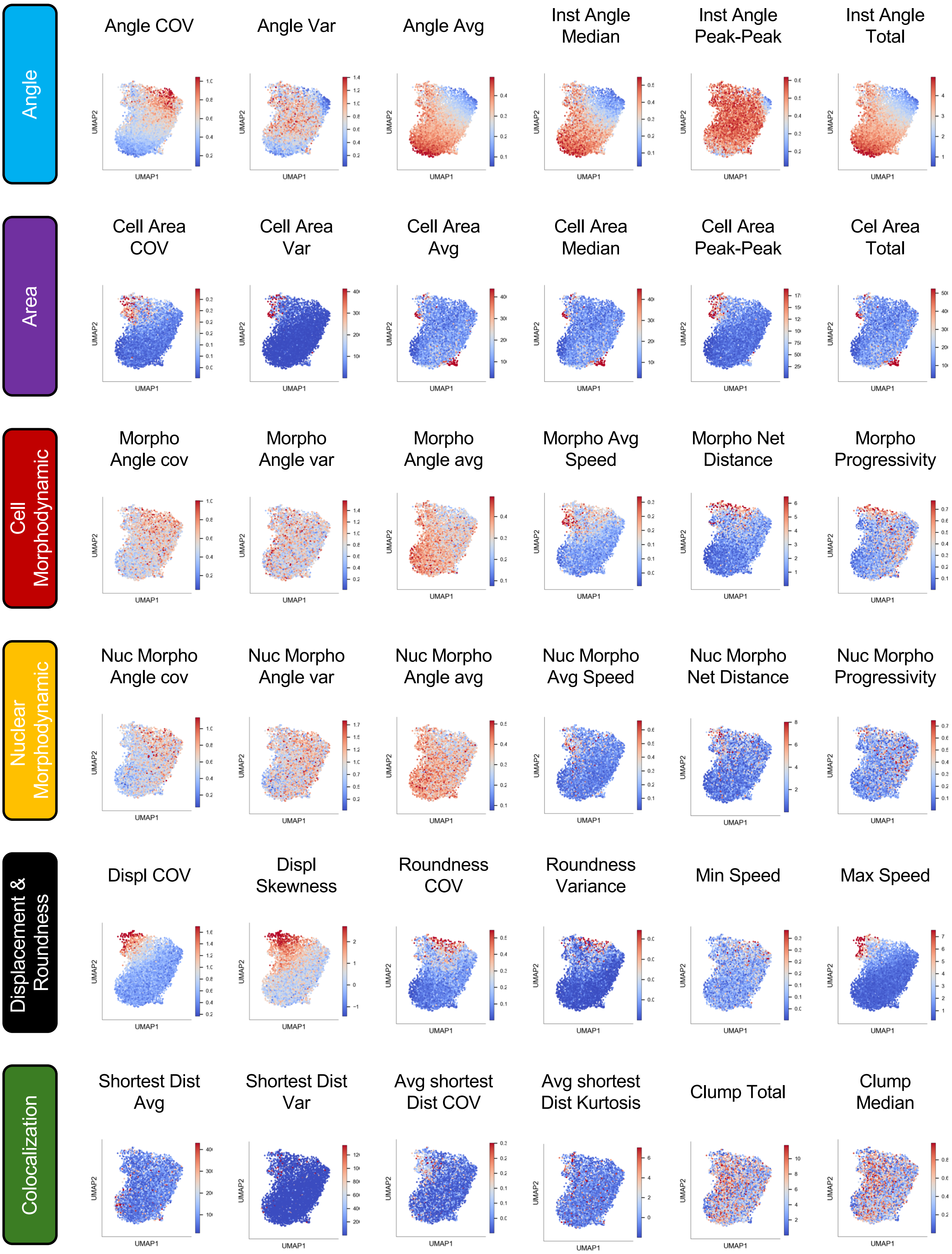

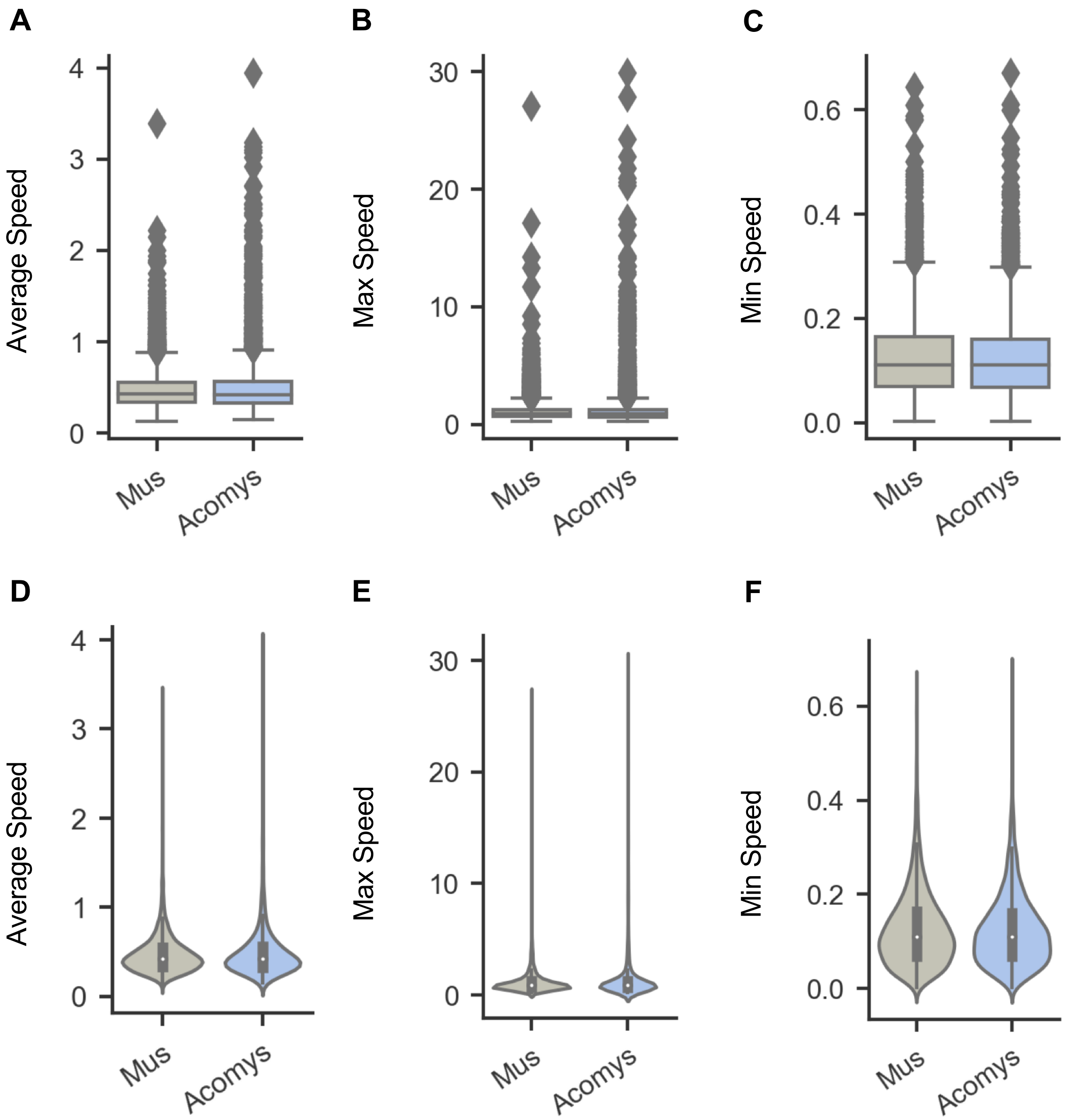
